# A chemically defined medium to support the growth of food-relevant *Bacillus* species

**DOI:** 10.1101/2025.07.14.664712

**Authors:** Tessa S. Canoy, Emma S. Wiedenbein, Charlie H. McPhillips, Lene Jespersen, Henriette L. Røder, Dennis S. Nielsen

**Affiliations:** Department of Food Science, Faculty of Science, University of Copenhagen, Frederiks- berg, Denmark

**Keywords:** *Bacillus*, defined medium, microbial cultivation, metabolism, growth

## Abstract

The *Bacillus* genus contains many members with food significance, including the food-grade *Bacillus subtilis* clade often used in fermentations and the pathogenic *Bacillus cereus* clade. Chemically defined media for *Bacillus* species are crucial tools to allow detailed investigations of the influence of specific nutrients on growth and also improve reproducibility and consistency of experiments. Previous studies have focused on the development of defined media for single species, while the aim of this study was to develop a chemically defined medium that supports the growth of multiple food relevant *Bacillus* species. The new medium, Pafoba, was tested using two pathogenic strains of the *Bacillus cereus* clade and eleven strains of the *Bacillus subtilis* clade representing seven different clade members. All thirteen *Bacillus* strains were able to grow on Pafoba, of which ten displayed a similar or higher maximum OD_600_ on Pafoba medium compared to rich medium (Brain Heart Infusion broth). Detailed analysis revealed a biotin requirement for *Bacillus subtilis* strain PRO64, and the necessity of including essential amino acids for *Bacillus weihenstephanensis* and *Bacillus cereus* strains. In conclusion, the chemically defined Pafoba medium provides a controlled and reproducible growth environment for fundamental studies and is suitable for detection and enumeration of a broad range of *Bacillus* spp. related to food processing and safety.

**Importance:** *Bacillus* species are important in both food fermentation and food safety. While members of the *Bacillus subtilis* clade are used in the production of fermented foods, those in the *Bacillus cereus* group are associated with foodborne illness. This study presents a chemically defined medium that supports the growth of multiple food-relevant *Bacillus* species, enabling precise control over nutrient composition. Knowledge of *Bacillus cereus* metabolism under defined conditions is essential to support efforts in food safety, risk assessment, and the development of targeted intervention strategies. Likewise, an understanding of *Bacillus subtilis* metabolism under defined conditions is essential to optimize fermentation processes, improve product consistency, and enhance functional food development. By providing a standardized and reproducible growth environment, the medium developed in this study will facilitate research that advances both microbial food safety and the controlled use of beneficial *Bacillus* strains in food production.

## Introduction

The *Bacillus* genus is a diverse group of rod-shaped Gram-positive sporeformers and contains two major clades: the *subtilis* clade, which includes 27 species, and the *cereus* clade, comprising 20 species (1). Taxonomically, these clades are grouped together based on a few shared phenotypic traits, including the ability to form endospores and grow aerobically. Advances in genomic and phylogenomic analyses have revealed that the genus *Bacillus* is highly polyphyletic, meaning that it includes species that are not all closely related in evolutionary terms (1, 2). Nevertheless, both *B. subtilis* and *B. cereus* groups have relevance to food production, either in the context of food fermentation and bioprotection (*Bacillus subtilis*) or for food safety (*Bacillus cereus*).

*Bacillus* (*B*.) *subtilis* is widely used as a model organism in laboratory research and is known for its diverse enzyme profile production (3). Multiple species belonging to the *B. subtilis* clade, including *B. licheniformis, B. amyloliquefaciens, B. pumilus* and *B. subtilis*, play essential roles in traditional fermented Asian and African foods (4, 5). *B. subtilis* inhabits diverse ecological niches, and foodrelevant species have been isolated from a wide range of sources, including foods, agricultural products and environmental reservoirs like soil, water, and intestinal tracts of animals (6). *Bacillus* strains belonging to the *B. subtilis* group have potential to transform plant ingredients into sustainable protein sources, through enhancement of nutritional value and flavor generation (7) as well as having bioprotective properties (8). At the same time, the presence of *B. subtilis* is not always desired, since it can spoil foods like bread (9) and dairy products (10) through its contribution to ropiness and offflavors.

*Bacillus cereus* on the other hand, is notable for its role in foodborne illnesses. The *Bacillus cereus* sensu lato group comprises a genetically close yet ecologically diverse clade of species, including *B. cereus* sensu stricto, *B. thuringiensis, B. anthracis, B. paranthracis* and *B. weihenstephanensis* (11). *B. cereus* can cause two types of gastrointestinal diseases, based on emetic toxin production (cereulide), which causes vomiting, and enterotoxin production, which causes diarrhea (12). *B. cereus* has been associated with many different types of foods, including cereals, beans, vegetables and dairy products (13). The spore-forming abilities of *B. cereus* complicates its effective control in food processing environments, since spores are resistant to many detergents (14) as well as physical treatments such as mild heat (pasteurization), boiling, drying and UV radiation (15).

To cultivate *Bacillus* species, rich media like Brain Heart Infusion Broth (BHI) or Tryptic Soy Broth (TSB) are typically used. However, these complex media are composed of undefined components, like meat extract and tryptone, that can vary in composition from batch to batch. Complex rich media are therefore unsuitable to provide a controlled environment for experimental studies. A chemically defined growth medium consists of pure chemicals with known concentrations. A defined medium has three important advantages: (1) it offers a reproducible growth environment, which minimizes variability, enhances the reliability of experimental results and facilitates direct comparisons between strains, (2) it allows for the precise control of nutritional variables, which can be used to study effects on biological processes, such as cell growth, sporulation and gene regulation and, (3) it eases comparison of results between different laboratories and studies.

The composition of defined media used for *Bacillus* species ranges from a few ingredients (minimal medium) to more elaborate ingredient lists (Table 1). A common factor in all *Bacillus* defined media is the use of mineral salts, such as dipotassium phosphate and magnesium sulphate. Typical carbon sources for *Bacillus* cultivation include glucose, maltose, L-sorbose, or sucrose, while the most common nitrogen sources are ammonium salts and amino acids (Table 1). Both calcium and manganese are essential for sporulation in *Bacillus* species (16–18), although the specific effect of calcium supplementation on sporulation rates may be dependent on the exact *Bacillus* species (19). Thiamine (vitamin B1) is added in some media for *Bacillus subtilis* (16, 20), while both thiamine and biotin (vitamin H) are added to media specifically designed for *Bacillus pumilus* (21, 22).

**Table 1:**
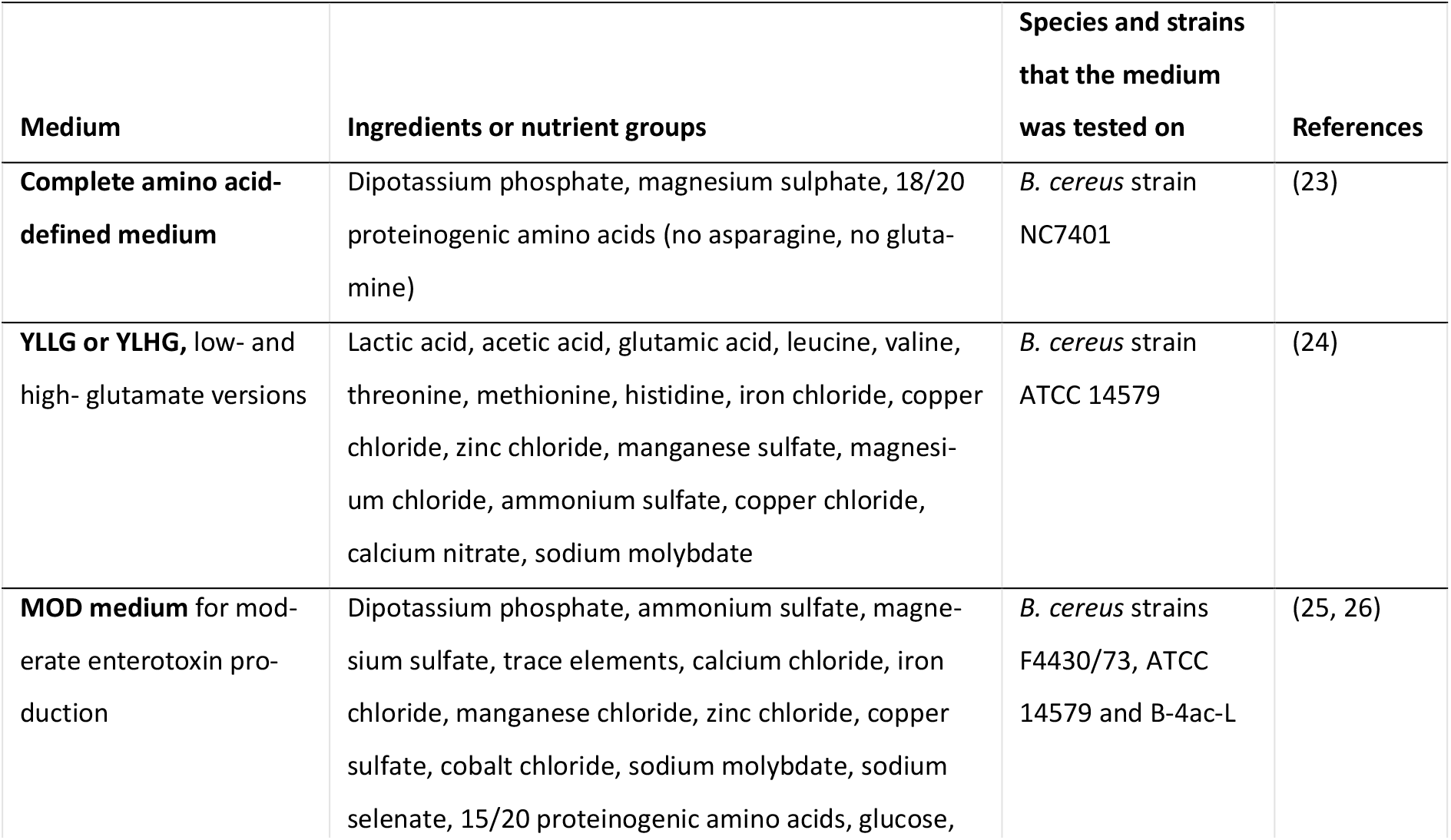
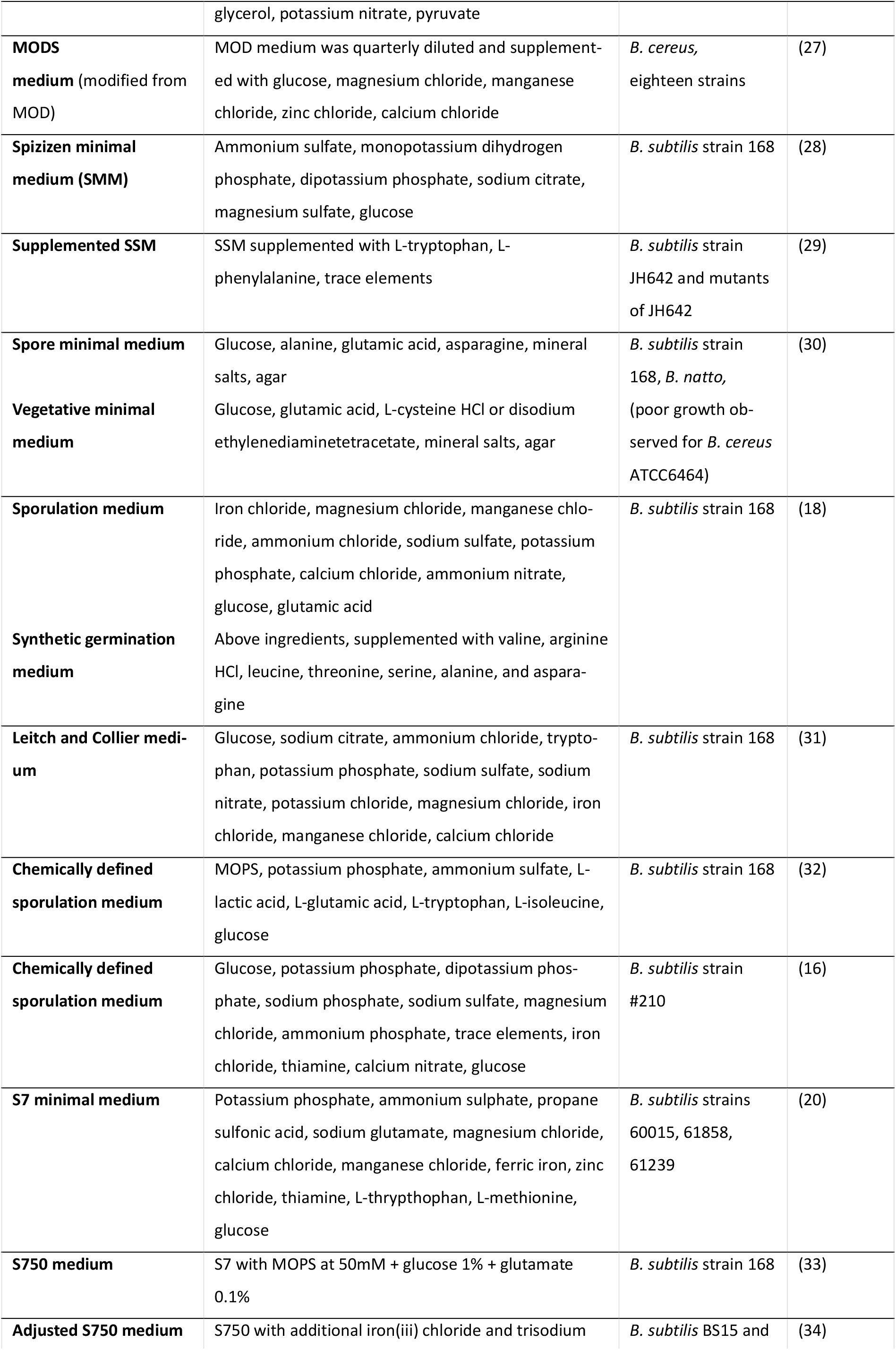
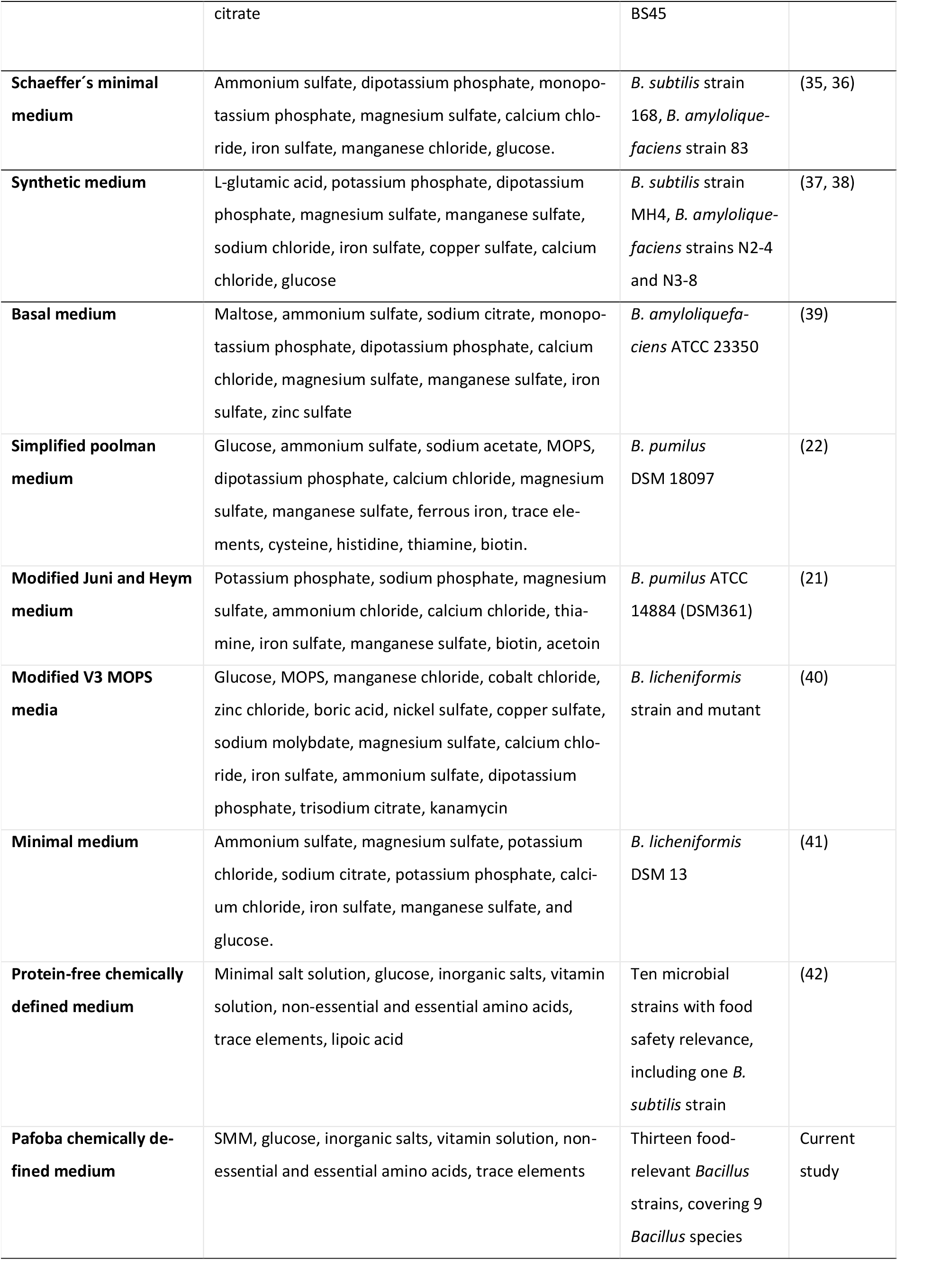
A selection of chemically defined media used for cultivation of *Bacillus* species from the *Bacillus cereus* and *Bacillus subtilis* clades. Abbreviation: MOPS morpholinepropanesulfonic acid.

A commonly used defined medium for *B. subtilis* cultivation is the Spizizen minimal medium (28). Spizizen minimal medium is frequently modified with additional components such as specific amino acids (e.g., tryptophan, phenylalanine, glutamate, alanine), trace elements, sodium chloride, or increased glucose concentrations (29, 43, 44). These modifications indicate that different *B. subtilis* strains can exhibit distinct nutritional requirements. This strain-dependent variability underscores the need to develop and validate new chemically defined media that are broadly applicable across diverse *Bacillus* isolates. Many known chemically defined media are tailored to individual strains or species only (Table 1), although a few exceptions exist. Schaeffer’s minimal medium (35) and the minimal medium by Jamil et al. (2007) (37) were originally developed for *Bacillus subtilis* but have also been adopted for cultivating *B. amyloliquefaciens* (36, 38). Furthermore, the broad range chemically defined medium developed by Wang, Greenwood and Klein (2021) (42) supports the cultivation of a variety of foodborne pathogens and spoilage organisms. This medium is particularly noteworthy as it supported growth of all ten tested strains at levels comparable to those observed in a rich medium such as TSB, while remaining a fully chemically defined medium. However, this medium was not optimized for the cultivation of *Bacillus* spp. and was only tested on one *Bacillus subtilis* strain.

Commonly, defined media have just been tested on the *Bacillus* type strain, *Bacillus subtilis* 168 (DSM10). Despite being widely used, the type-strain does not reflect the broad metabolic and genetic diversity within neither the *subtilis* clade nor the *cereus* clade. The aim of this study was therefore to develop a chemically defined medium capable of supporting the growth of multiple *Bacillus* species, including both food-relevant and pathogenic strains, to allow for more reproducible comparative studies across the *Bacillus* genus.

## Materials and methods

### Preparation of chemically defined media

Three chemically defined media were prepared for this study: Spizizen minimal medium, as originally described by Anagnostopoulos & Spizizen (1961) (28); a modified version of the medium developed by Wang, Greenwood and Klein (2021) (42), hereafter referred to as WGK medium; and a further developed formulation named Pafoba, optimized for the growth of PAthogenic and FOod-grade *Bacillus* species. The compositions of the WGK and Pafoba media are provided in Table 3.

To prepare the WGK and Pafoba media, the base ingredients were dissolved in demineralized water at concentrations higher than their final working concentrations, such that the complete medium would reach the values listed in Table 3. The base mixture along with 50x glucose and 100x calcium chloride stock solutions were autoclaved at 121 °C for 20 minutes. Inorganic salts solution (100x) was sterilized through a 0.22 µM filter, stored at 4°C and used within two months after preparation. Biotin stock solution (100x) was prepared fresh. The final medium was prepared by mixing the autoclaved base solution with appropriate volumes of the stock solutions glucose (50x), calcium chloride (100x), Gibco MEM amino acids solution (50x), Gibco MEM non-essential amino acids solution (100x), Gibco MEM vitamin solution (100x), biotin (200x), and Trace Metal mix solution (1000x). The complete medium was vacuum filter-sterilized through a 0.2 µM filter (Nalgene, Thermo Scientific). During preparation, exposure to light was minimized to preserve the light-sensitive biotin. The pH of the completed defined medium (6.7 ± 0.1 after filter-sterilization) was not adjusted. The medium containers were stored at -60°C and used within three weeks after preparation.

### Bacterial strains

The Spizizen minimal medium was tested on 37 non-pathogenic *Bacillus* strains, including 19 *B. subtilis*, 8 *B. velezensis*, 3 *B. pumilus*, 3 *B. paralicheniformis*, 3 *B. licheniformis*, and 1 *B. amyloliquefaciens* strains, all described by Wiedenbein et al. (2023) (45). To test the WGK and Pafoba media, thirteen *Bacillus* strains, including two pathogenic and eleven non-pathogenic strains (Table 2), were used. Seven non-pathogenic *Bacillus* strains were bought from the German Collection of Microorganisms and Cell Cultures (DSMZ, Germany). Four *Bacillus* strains were selected that had previously been isolated from food fermentations in Ghana and Burkina Faso (4, 45, 46). From the *B. cereus* sensu lato group, one *Bacillus weihenstephanensis* and one *B. cereus* sensu stricto were included. Pathogenic strain F4810/72 was previously isolated from a foodborne outbreak and identified as *B. cereus* (47). *B. weihenstephanensis* MC67 was previously isolated from sandy soil (48).

**Table 2:** Composition of the chemically defined media WGK and Pafoba used for the cultivation of *Bacillus* species.

| Compound | WGK concentration (mM) | Pafoba concentration (mM) | Supplier |
| --- | --- | --- | --- |
| **Base ingredients** |  |  |  |
| KH<sub>2</sub>PO<sub>4</sub> | 22 | 44 | Merck |
| K<sub>2</sub>HPO<sub>4</sub> | - | 80 | Merck |
| Na<sub>2</sub>HPO<sub>4</sub> | 90 | - | Merck |
| Na<sub>3</sub>C<sub>6</sub>H<sub>5</sub>O<sub>7</sub> · 2H<sub>2</sub>O | - | 3.4 | Merck |
| NaCl | 8.56 | - | Sigma-Aldrich |
| (NH<sub>4</sub>)<sub>2</sub>SO<sub>4</sub> | - | 15 | Sigma-Aldrich |
| NH<sub>4</sub>Cl | 1.87 | - | Sigma-Aldrich |
| **D-Glucose** | 22 | 28 | Sigma-Aldrich |
| **Inorganic salts** |  |  |  |
| MgSO<sub>4</sub> x H<sub>2</sub>O | 2 | 2 | VWR |
| FeSO<sub>4</sub> x 7 H<sub>2</sub>O | 0.04 | 0.15 | Sigma-Aldrich |
| MnSO<sub>4</sub> x H<sub>2</sub>O | 0.05 | 0.05 | Merck |
| **Calcium chloride (CaCl<sub>2</sub>·2H<sub>2</sub>O)** | 1.5 | 1.5 | Merck |
| **Trace Metal Mix A5 with Co** |  |  | Sigma-Aldrich |
| H<sub>3</sub>BO<sub>3</sub> | 0.046 | 0.046 |  |
| MnCl<sub>2</sub>·4H<sub>2</sub>O | 0.009 | 0.009 |  |
| ZnSO<sub>4</sub>·7H<sub>2</sub>O | 0.00077 | 0.00077 |  |
| NaMoO<sub>4</sub>·2H<sub>2</sub>O | 0.0016 | 0.0016 |  |
| CuSO<sub>4</sub>·5H<sub>2</sub>O | 0.00032 | 0.00032 |  |
| Co(NO<sub>3</sub>)<sub>2</sub>·6H<sub>2</sub>O | 0.00017 | 0.00017 |  |
| **Gibco MEM amino acids solution** |  |  | ThermoFisher |
| L-Arginine hydrochloride | 0.6 | 0.6 |  |
| L-Cysteine | 0.1 | 0.1 |  |
| L-Histidine hydrochloride | 0.2 | 0.2 |  |
| L-Isoleucine | 0.4 | 0.4 |  |
| L-Leucine | 0.4 | 0.4 |  |
| L-Lysine hydrochloride | 0.4 | 0.4 |  |
| L-Methionine | 0.1 | 0.1 |  |
| L-Phenylalanine | 0.2 | 0.2 |  |
| L-Threonine | 0.4 | 0.4 |  |
| L-Tryptophan | 0.05 | 0.05 |  |
| L-Tyrosine | 0.2 | 0.2 |  |
| L-Valine | 0.4 | 0.4 |  |
| **Biotin** | 2.44 | 2.44 | Sigma-Aldrich |
| **Gibco MEM vitamin solution** |  |  | ThermoFisher |
| Choline chloride | 0.007 | 0.007 |  |
| D-Calcium pantothenate | 0.002 | 0.002 |  |
| Folic acid | 0.002 | 0.002 |  |
| Niacinamide | 0.008 | 0.008 |  |
| Pyridoxal hydrochloride | 0.005 | 0.005 |  |
| Riboflavin | $2.6 \times 10^{-5}$ | $2.6 \times 10^{-5}$ | |
| Thiamine hydrochloride | 0.003 | 0.003 |  |
| DL-Inositol | 0.01 | 0.01 |  |
| NaCl | 1.45 | 1.45 |  |
| **Gibco MEM non-essential** |  |  | ThermoFisher |
| Glycine | 0.1 | 0.1 |  |
| L-Alanine | 0.1 | 0.1 |  |
| L-Asparagine | 0.1 | 0.1 |  |
| L-Aspartic acid | 0.1 | 0.1 |  |
| L-Glutamic acid | 0.1 | 0.1 |  |
| L-Proline | 0.1 | 0.1 |  |
| L-Serine | 0.1 | 0.1 |  |

**Table 3:** Characteristics of *Bacillus* strains included in the present study. T refers to type strain.

| Strains | Role in food | Isolation source, country of origin |
| --- | --- | --- |
| *Bacillus cereus* F4810/72 | Pathogen (emetic) | Foodborne outbreak, UK |
| *Bacillus weihenstephanensis* MC67 | Pathogen (psychrotolerant) | Sandy soil, Denmark |
| *Bacillus subtilis* PRO64 | Fermentation | Kantong (Ghana) |
| *Bacillus subtilis* PRO122 | Fermentation | Maari (Burkina Faso) |
| *Bacillus subtilis* DSM21393 | Fermentation | Potato tubers Tatarstan (Russian Federation) |
| *Bacillus subtilis* DSM10<sup>T</sup> | Fermentation | Unknown ("Marburg") |
| *Bacillus amyloliquefaciens* PRO40 | Fermentation | Kantong (Ghana) |
| *Bacillus velezensis* PRO96 | Fermentation | Kantong (Ghana) |
| *Bacillus licheniformis* DSM26543 | Fermentation | Lung of mouse experimental animal, Plön (Germany) |
| *Bacillus licheniformis* DSM1969 | Fermentation | Soil |
| *Bacillus licheniformis* DSM13<sup>T</sup> | Fermentation | Unknown |
| *Bacillus pumilus* DSM1794 | Fermentation | Paperboard from a moldy building, Mikkeli (Finland) |
| *Bacillus pumilus* DSM13835 | Fermentation | Indoor air of water damaged building (Finland) |

### Cultivation and OD_600_ measurement

Strains were pre-cultured in 10 mL Brain Heart Infusion (BHI) broth (HiMedia, Germany) twice overnight at 30°C and 200 rpm. After the second overnight culture, cells were harvested through centrifugation (5000g, 10 min). To avoid carryover from BHI medium, the cells were washed with salt solution (0.9% NaCl) twice. The washed cells were suspended in 10 mL salt solution, adjusted to an OD_600_ of 0.2 before being used as inoculum.

Each well of a 96-well microplate was filled with 2 µL inoculum and 198 µL medium and sealed with an oxygen-permeable film (Breath-Easy sealing membrane, Sigma Aldrich). The growth of each strain on the chemically defined media was monitored by measuring the optical density at 600 nm (OD_600_). OD_600_ was measured every hour with a Biotek ELx808 Absorbance Microplate Reader (Biotek, USA) during 48h at 30°C, with high shake speed prior to each measurement. For all experiments and strains, three biological replicates and two technical replicates were used. Wells with non-inoculated medium were used as negative control and to adjust for background absorbance. The growth of all thirteen *Bacillus* strains (Table 2) was monitored during cultivation on WGK, Pafoba, and BHI, with BHI as a rich medium control.

### Nutrient substitutions and enrichments in WGK medium

To evaluate the contribution of specific nutrients of the Pafoba medium to *Bacillus* growth, three strains were selected for cultivation in modified WGK media: *Bacillus subtilis* PRO64, *Bacillus subtilis* DSM21393, and *Bacillus licheniformis* DSM13 ^T^. Five variants of WGK medium were prepared, each containing a single nutrient substitution or enrichment to reflect the concentrations found in the Pafoba medium (Table 4). These included two nutrient substitutions (ammonium chloride replaced with ammonium sulfate, sodium chloride replaced with trisodium citrate), and three nutrient enrichments (glucose, iron sulfate, KH_2_PO_4_). Growth on the five adjusted WGK media was compared with the growth on the original WGK and Pafoba medium. The difference in composition between K_2_HPO_4_ (80 mM, Pafoba) and Na_2_HPO_4_ (90 mM, WGK) was not assessed.

**Table 4:** Differences in composition between the Pafoba and WGK chemically defined media, and the five adjusted WGK media each representing single nutrient substitutions or enrichments, to test for the effect of individual nutrients in the Pafoba medium on the growth of *Bacillus* strains.

| WGK medium | WGK media with single nutrient substitutions or enrichments | Pafoba medium |
| --- | --- | --- |
| Ammonium chloride (1.87 mM) | 1. WGK medium with ammonium sulfate (instead of ammonium chloride) | Ammonium sulfate (15 mM) |
| Sodium chloride (8.56 mM) | 2. WGK medium with trisodium citrate (instead of sodium chloride) | Trisodium citrate (3.4 mM) |
| Less iron sulfate (0.04 mM) | 3. WGK medium with 3.7x more iron sulfate | More iron sulfate (0.15 mM) |
| Less glucose (22 mM) | 4. WGK medium with 1.3x more glucose | More glucose (28 mM) |
| Less $\text{KH}_2\text{PO}_4$ (22 mM) | 5. WGK medium with 2x more potassium phosphate | More potassium dihydrogen phosphate (44 mM) |

### Nutrient-deprived Pafoba media to test for auxotrophies

To get a better understanding of the nutrient auxotrophies of the individual strains and investigate the possibilities for simplifying medium composition, the growth of *Bacillus* strains on variations of the Pafoba medium without specific nutrients was monitored. Pafoba medium was prepared, while single nutrient or nutrient groups were omitted, resulting in five nutrient-deprived versions of Pafoba media: (1) Pafoba without biotin, (2) Pafoba without MEM vitamins solution, (3) Pafoba without MEM essential amino acids, (4) Pafoba without MEM non-essential amino acids, and (5) Pafoba without Trace element mix. The maximum OD_600_ during 48 hours of growth as well as the time to reach the maximum OD_600_ on each of the media was determined. Deprived nutrient groups were classified as required (Max OD_600_ deprived medium <25% than Max OD_600_ Pafoba), stimulatory (25 % Max OD_600_ Pafoba ≤ Max OD_600_ deprived medium ≤ 75 % Max OD_600_ Pafoba), or negligible (Max OD_600_ D600 deprived medium >75 % Max OD_600_ Pafoba).

### Data analysis

Data analysis and statistical analysis were performed using R version 4.2.1 (49). OD_600_ was corrected for background medium. Average maximum OD600 (Max OD_600_) and time to reach MaxOD600 was calculated based on biological replicates and technical replicates, while standard deviation was based on biological replicates. Max OD_600_ and time to reach Max OD_600_ were compared between Pafoba, WGK and BHI media using ANOVA, with Dunn with Bonferroni as post-hoc tests. To evaluate the effect of nutrient substitutions and enrichments on *Bacillus* growth, Max OD_600_ was compared between Pafoba, WGK and the five adjusted WGK media (Table 4) using ANOVA. Post-hoc Dunn with Bonferroni was performed, using WGK and Pafoba as control conditions for comparison with each adjusted WGK medium. A significance level of 0.05 was used for all statistical tests.

## Results

### Development of the Pafoba medium

Spizizen minimal medium was initially evaluated for its suitability to support the growth of a variety of *Bacillus* strains. We observed that 20 out of 37 non-pathogenic *Bacillus* strains described by Wiedenbein et al. (2023) (45) grew on the Spizizen medium. However, growth was poor, with optical densities (OD600) typically reaching only 0.1-0.2 after 48 hours (data not shown). Given this limited performance, we turned our attention to the medium described by Wang, Greenwood and Klein (2021) (42), which had previously been shown to support sustained growth of various food-relevant microbes at levels comparable to those seen in TSB medium. Building on this formulation, we developed an adapted version tailored for *Bacillus* spp., referred to as WGK medium. This modified medium retained most components of the study by Wang, Greenwood and Klein (2021) (42), but with two adjustments: lipoic acid was excluded, as preliminary screening showed it was not required for *Bacillus* growth (Supplementary Figure S1), and the biotin concentration was increased to improve for the growth of *B. pumilus* DSM13835.

Despite these improvements, WGK medium did not support robust growth of all *Bacillus* strains. Notably, certain *B. subtilis* and *B. licheniformis* strains exhibited poor growth on WGK medium (Figure 1A). To address this limitation, we developed a second version, named Pafoba medium. The Pafoba medium was developed by substituting the M9 minimal salts solution of the WGK medium with salts of the Spizizen minimal medium, consisting of ammonium sulfate, sodium citrate, monopotassium dihydrogen phosphate, and dipotassium phosphate. Additionally, we increased the concentrations of glucose and potassium dihydrogen phosphate, similarly to the Spizizen minimum medium, and incorporated a higher iron sulfate concentration (41 mg/L), based on the work of Coltin et al. (2022) (50) who optimized trace element requirements of *Priestia megaterium*, formerly *Bacillus megaterium*.

### Growth measurement on WGK, Pafoba and BHI media

To evaluate the performance of the newly developed Pafoba medium, the growth of thirteen *Bacillus* strains was monitored over 48h on WGK, Pafoba and BHI media (Figure 1A). Growth performance was assessed by comparing maximum OD_600_ and time to reach maximum OD_600_ (Figure 1B). Compared to WGK medium, Pafoba supported equal or superior growth for all thirteen *Bacillus* strains. For eight *Bacillus* strains, growth on Pafoba resulted in a higher maximum OD_600_ than on WGK, of which three *B. licheniformis* strains (DSM13 type strain, DSM1969, DSM26543), three *B. subtilis* strains (DSM21393, PRO64, PRO122), *B. pumilus* DSM13835, and *B. cereus* F4810/72. While *B. velezensis* PRO96 reached a similar maximum OD_600_ in Pafoba and WGK, it reached max OD_600_ earlier on WGK (18 ± 1h) than on Pafoba (27 ± 2h).

**Figure 1:**
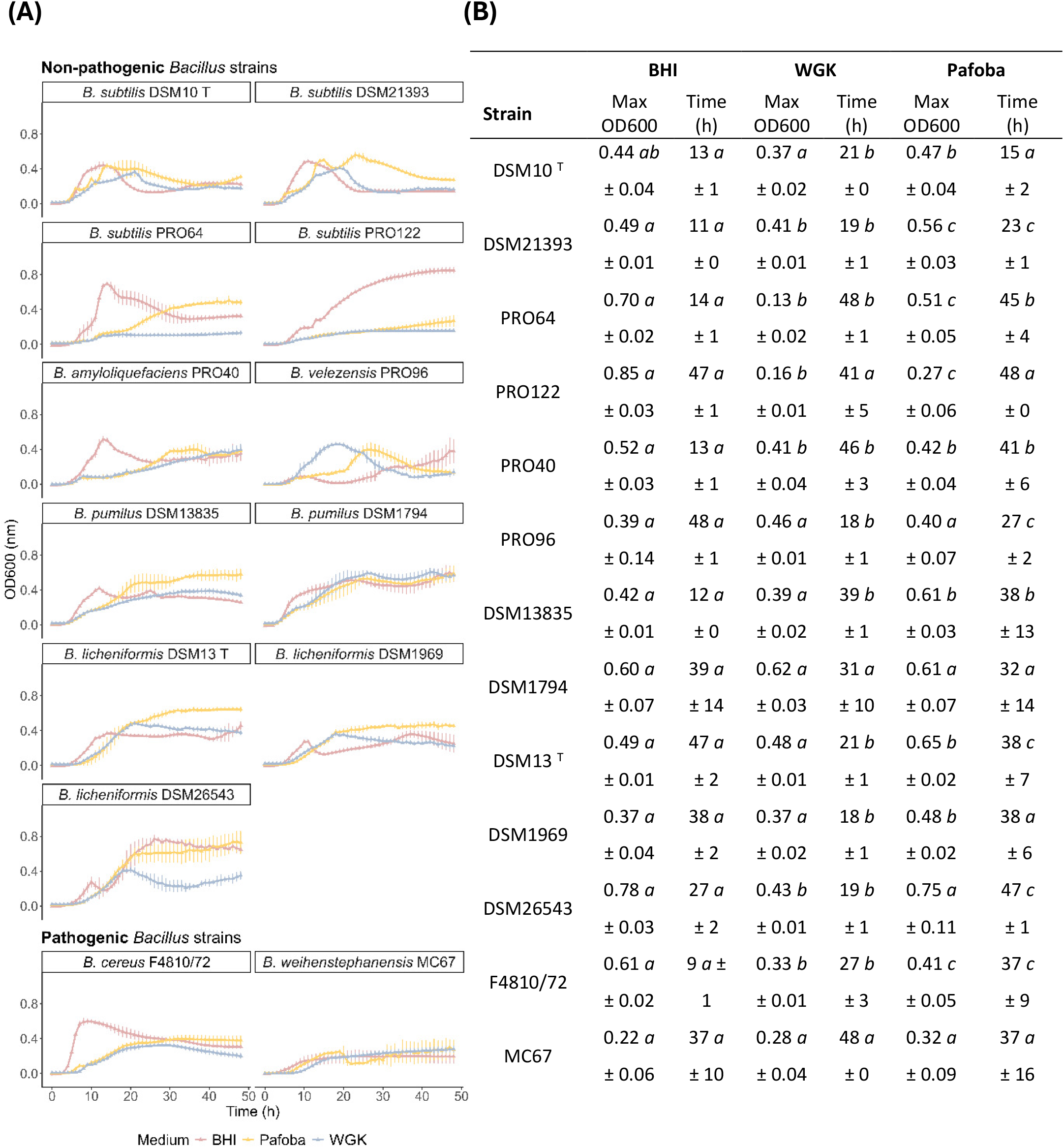
Comparison of growth (OD600) by *Bacillus* strains on Pafoba and WGK chemically defined media and BHI (rich medium) (A) Growth for 48 hours (B) Maximum OD600 and time (h) to reach maximum OD600 during 48h cultivation. Means denoted by a different letter indicate significant differences between treatments (p < 0.05).

Growth in the nutrient-rich BHI medium was generally faster than in Pafoba medium, although the maximum OD_600_ values were similar in most cases. Six *Bacillus* strains reached max OD_600_ earlier on BHI than on Pafoba, including two *B. subtilis* strains (DSM21393, PRO64), *B. licheniformis* DSM26543, *B. amyloliquefaciens* PPRO40, *B. pumilus* DSM13835, and *B. cereus* F4810/72. For four strains (PRO122, DSM1794, MC67, DSM10), time to reach max OD_600_ was similar between BHI and Pafoba. Notably, ten out of thirteen *Bacillus* strains displayed a similar or higher maximum OD_600_ in Pafoba medium compared to BHI (Figure 1). Even among the three strains that grew better on BHI medium, two *Bacillus* strains (*B. subtilis* PRO122, *B. cereus* F4810/72) showed better growth on Pafoba than on WGK medium. Overall, the Pafoba medium supported robust growth across a broad range of *Bacillus* strains, outperformed the WGK medium for eight strains, and approached the growth performance of the rich BHI medium for most strains.

### Nutrient substitutions and enrichments in WGK medium

Three *Bacillus* strains that displayed better growth on Pafoba than on WGK medium (Figure 1) were selected for further analysis: *Bacillus subtilis* PRO64, *Bacillus subtilis* DSM21393, and *Bacillus licheniformis* DSM13 ^T^. To better understand the distinct effect of the components of the WGK and Pafoba medium on *Bacillus* growth, five adjusted WGK media were prepared with each medium reflecting a single nutrient substitution or enrichment, including two media with nutrient substitutions (ammonium sulfate/ammonium chloride and sodium citrate/sodium chloride) and three media with nutrient enrichments (potassium dihydrogen phosphate, iron sulfate, glucose) (Table 4). The growth of the three *Bacillus* strains on Pafoba, WGK and adjusted WGK media is shown in Figure 2.

**Figure 2:**
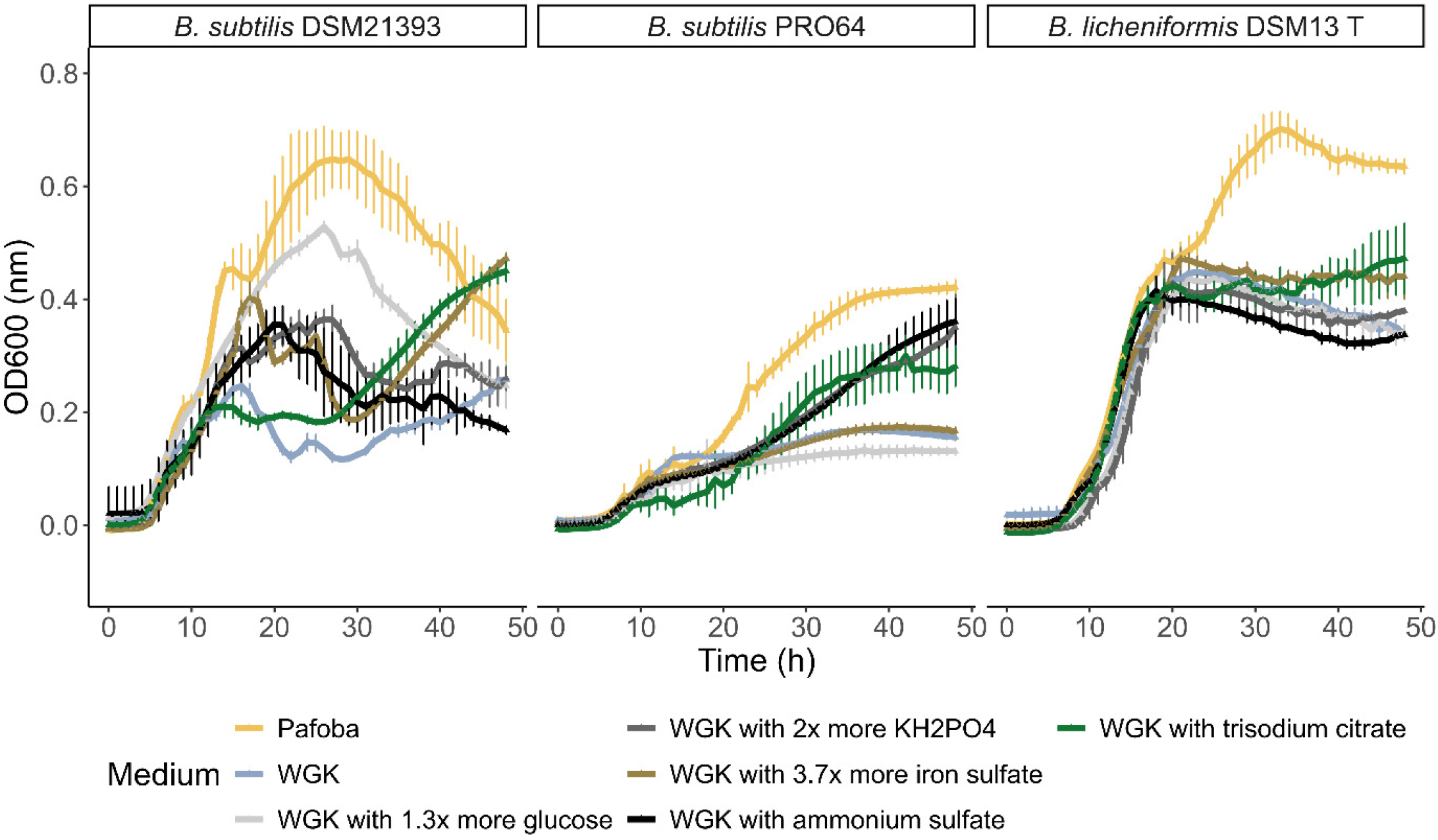
Effect of enriching or substituting nutrients of the WGK chemically defined medium on the growth (OD600) of *Bacillus* strains, including the Pafoba medium as the optimal growth medium. See the main text for statistical results.

All three *Bacillus* strains exhibited the highest growth in the Pafoba medium compared to the original WGK medium and the five modified WGK formulations (Figure 2). Nutrient replacement and enrichment of the WGK medium exerted strain-specific effects on *Bacillus* growth. *B. licheniformis* DSM 13 ^T^ grew significantly to significantly higher OD_600_ on the Pafoba medium (0.70 ± 0.03) than on the WGK medium (0.45 ± 0.01). None of the adjusted WGK media significantly (p<0.05) increased the max OD_600_ of *B. licheniformis* DSM13 ^T^ compared to the non-adjusted WGK medium. For *B. licheniformis* DSM13 ^T^, substitution of sodium chloride by trisodium citrate significantly increased the time to reach Max OD_600_ (41 ± 10h) compared to WGK (23 ± 1.5h). In contrast, for *B. subtilis* DSM21393 all the adjusted WGK media resulted in a higher max OD_600_ than the WGK medium. Among the adjusted WGK media, the medium with enriched glucose content resulted in the highest max OD_600_ (0.53 ± 0.01), although still lower than the Pafoba medium (0.66 ± 0.06) (p = 1.3*10^−4^). For *B. subtilis* PRO64, three modifications significantly promoted growth compared to the WGK medium: substitution of sodium chloride (8.6 mM) by trisodium citrate (3.4 mM) (p = 1.7*10^−4^), substitution of ammonium chloride by ammonium sulfate (p = 6.3*10^−6^), and a twofold enrichment of KH2PO4 (p = 1.0*10^−5^). The remaining modifications (more glucose, more iron sulfate) had no significant impact on *B. subtilis* PRO64 growth compared to the original WGK medium.

### Removal of nutrient groups from Pafoba medium to test for auxotrophies

Five versions of Pafoba medium with specific nutrients or nutrients groups omitted were developed to assess their individual impact on *Bacillus* growth: (1) biotin, (2) trace metals, (3) vitamins, (4) essential amino acids, and (5) non-essential amino acids. From all thirteen *Bacillus* strains tested, only the *B. subtilis* DSM 10 ^T^ was unaffected by nutrient omissions from the five depleted media (Table 5). Growth of the remaining twelve *Bacillus* strains was affected by the absence of at least one or more nutrient groups. Supplementation with essential amino acids was the most influential nutrient group, as their addition to the Pafoba medium stimulated or was required for growth in nine out of the thirteen tested strains. The two pathogenic strains, *Bacillus cereus* F4810/72 and *Bacillus weihenstephanensis* MC67, were distinct from the remaining eleven food-grade *Bacillus* strains, as they required essential amino acids for growth. The second most influential nutrient group was nonessential amino acids, which stimulated growth in seven out of thirteen strains. Biotin was stimulatory for five out of thirteen strains, while biotin was required for growth of *B. subtilis* PRO64. Least important were the trace metals and vitamin mixture, which both stimulated growth of three *Bacillus* strains.

**Table 5:**
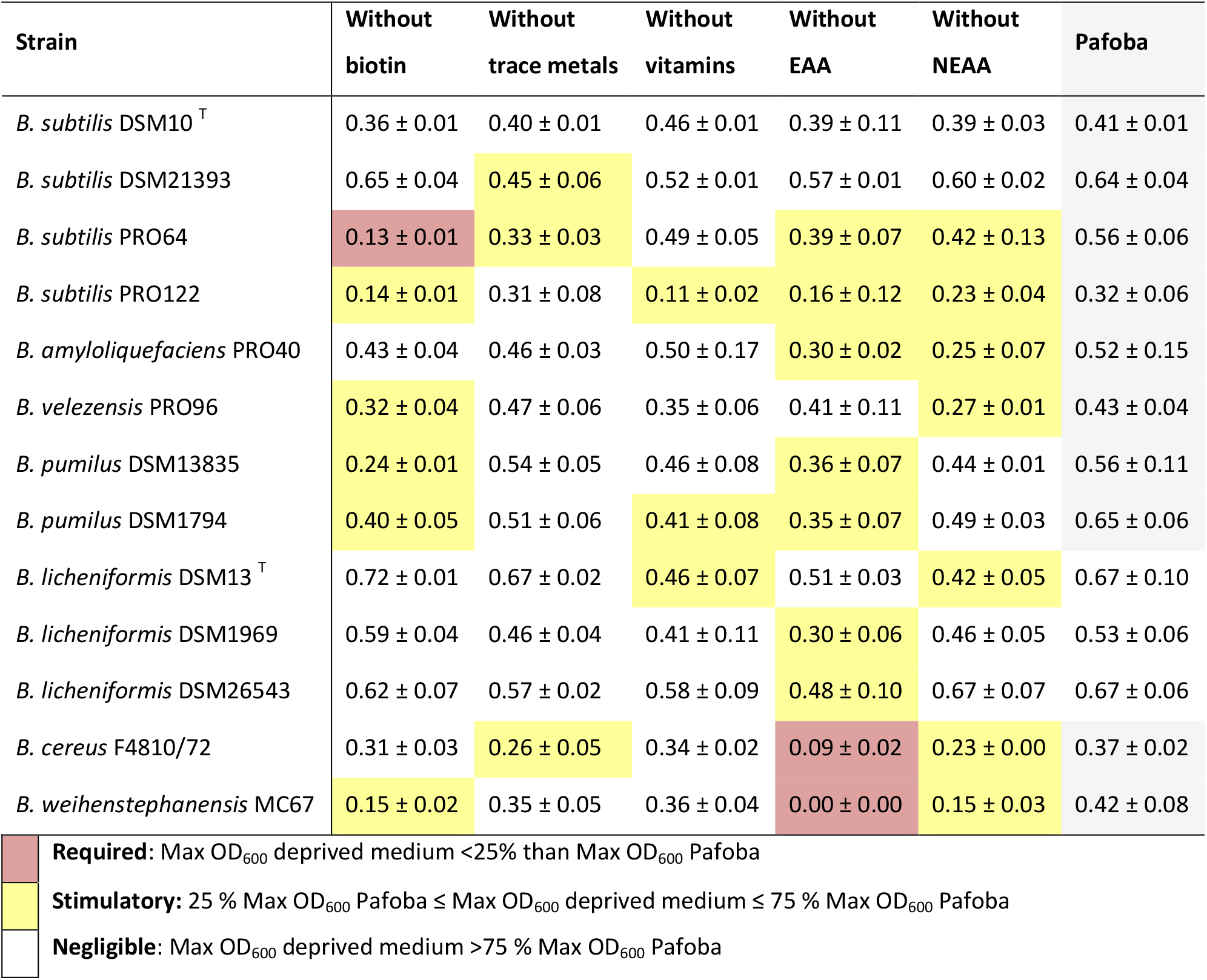
Effect of omitting specific nutrient groups from Pafoba medium on the maximum growth (maximum OD_600_) of *Bacillus* strains for 48 hours, with the complete Pafoba medium in the last column as a reference. Values represent the average of three biological replicates and two technical replicates and the standard deviation of three biological replicates. Cell colors are according to footnote. Abbreviations EAA: essential amino acids, NEAA: non-essential amino acids.

| Strain | Without<br>biotin | Without<br>trace metals | Without<br>vitamins | Without<br>EAA | Without<br>NEAA | Pafoba |
| --- | --- | --- | --- | --- | --- | --- |
| *B. subtilis* DSM10<sup>T</sup> | 0.36 ± 0.01 | 0.40 ± 0.01 | 0.46 ± 0.01 | 0.39 ± 0.11 | 0.39 ± 0.03 | 0.41 ± 0.01 |
| *B. subtilis* DSM21393 | 0.65 ± 0.04 | 0.45 ± 0.06 | 0.52 ± 0.01 | 0.57 ± 0.01 | 0.60 ± 0.02 | 0.64 ± 0.04 |
| *B. subtilis* PRO64 | 0.13 ± 0.01 | 0.33 ± 0.03 | 0.49 ± 0.05 | 0.39 ± 0.07 | 0.42 ± 0.13 | 0.56 ± 0.06 |
| *B. subtilis* PRO122 | 0.14 ± 0.01 | 0.31 ± 0.08 | 0.11 ± 0.02 | 0.16 ± 0.12 | 0.23 ± 0.04 | 0.32 ± 0.06 |
| *B. amyloliquefaciens* PRO40 | 0.43 ± 0.04 | 0.46 ± 0.03 | 0.50 ± 0.17 | 0.30 ± 0.02 | 0.25 ± 0.07 | 0.52 ± 0.15 |
| *B. velezensis* PRO96 | 0.32 ± 0.04 | 0.47 ± 0.06 | 0.35 ± 0.06 | 0.41 ± 0.11 | 0.27 ± 0.01 | 0.43 ± 0.04 |
| *B. pumilus* DSM13835 | 0.24 ± 0.01 | 0.54 ± 0.05 | 0.46 ± 0.08 | 0.36 ± 0.07 | 0.44 ± 0.01 | 0.56 ± 0.11 |
| *B. pumilus* DSM1794 | 0.40 ± 0.05 | 0.51 ± 0.06 | 0.41 ± 0.08 | 0.35 ± 0.07 | 0.49 ± 0.03 | 0.65 ± 0.06 |
| *B. licheniformis* DSM13<sup>T</sup> | 0.72 ± 0.01 | 0.67 ± 0.02 | 0.46 ± 0.07 | 0.51 ± 0.03 | 0.42 ± 0.05 | 0.67 ± 0.10 |
| *B. licheniformis* DSM1969 | 0.59 ± 0.04 | 0.46 ± 0.04 | 0.41 ± 0.11 | 0.30 ± 0.06 | 0.46 ± 0.05 | 0.53 ± 0.06 |
| *B. licheniformis* DSM26543 | 0.62 ± 0.07 | 0.57 ± 0.02 | 0.58 ± 0.09 | 0.48 ± 0.10 | 0.67 ± 0.07 | 0.67 ± 0.06 |
| *B. cereus* F4810/72 | 0.31 ± 0.03 | 0.26 ± 0.05 | 0.34 ± 0.02 | 0.09 ± 0.02 | 0.23 ± 0.00 | 0.37 ± 0.02 |
| *B. weihenstephanensis* MC67 | 0.15 ± 0.02 | 0.35 ± 0.05 | 0.36 ± 0.04 | 0.00 ± 0.00 | 0.15 ± 0.03 | 0.42 ± 0.08 |

|  |  |
| --- | --- |
|  | **Required:** Max OD<sub>600</sub> deprived medium <25% than Max OD<sub>600</sub> Pafoba |
|  | **Stimulatory:** 25 % Max OD<sub>600</sub> Pafoba ≤ Max OD<sub>600</sub> deprived medium ≤ 75 % Max OD<sub>600</sub> Pafoba |
|  | **Negligible:** Max OD<sub>600</sub> deprived medium >75 % Max OD<sub>600</sub> Pafoba |

## Discussion

Here we report the development of a new chemically defined medium, Pafoba, which supports the growth of multiple species of the *B. subtilis* and *B. cereus* clades, enabling direct and standardized comparisons between diverse *Bacillus* strains and studies. Out of thirteen *Bacillus* strains, ten strains displayed a similar or better growth performance in Pafoba medium compared to the complex rich medium BHI. Growth was generally faster on BHI compared to the defined media. All *Bacillus* strains exhibited either comparable growth (five strains) or superior growth (eight strains) on the newly formulated Pafoba medium compared to the previously developed WGK medium.

To further understand the improved growth performance of *Bacillus* on Pafoba compared to WGK, growth of one *B. licheniformis* and two *B. subtilis* strains were monitored on WGK, Pafoba and five adjusted WGK media that each contained a single nutrient substitution or enrichment. For the type strain *B. licheniformis* DSM13^T^, none of the adjusted WGK media significantly improved growth compared to the WGK medium. Three nutrient adjustments of the WGK medium improved the growth of two *B. subtilis* strains (PRO64 and DSM21393): i) an increased amount of potassium phosphate, ii) the substitution of ammonium chloride by ammonium sulfate, and iii) the substitution of sodium chloride by trisodium citrate. For *B. subtilis* DSM21393, higher amounts of glucose and iron sulfate additionally enhanced growth.

Enrichment by potassium phosphate could have increased growth, as phosphate is a critical component of cellular molecules, like ATP, peptidoglycan and phospholipids (51). In addition, potassium phosphate supplementation has been shown to increase the heat resistance of *B. cereus* and *B. subtilis* spores (52, 53). Substitution of ammonium chloride by ammonium sulfate likely contributed to *Bacillus* growth, as sulfate serves as a primary source of sulfur and influences broader metabolic activities of *B. subtilis*, like amino acid synthesis, carbon metabolism and oxidative stress response (54). The substitution of sodium chloride by trisodium citrate also stimulated *B. subtilis* growth. Trisodium citrate is a chelator of divalent metal ions (e.g. Ca^2+^, Mg^2+^) and is a readily assimilable form of citrate, providing a carbon source that can be directly metabolized in the citric acid cycle, bypassing both glycolysis and pyruvate conversion (55). Trisodium citrate supplementation has been shown to increase the *B. licheniformis* production of poly-γ-glutamic acid, a highly viscous polymer responsible for the typical natto structure (56). These results highlight the importance of sulfate, potassium phosphate and carbon availability (trisodium citrate, glucose) for *Bacillus* metabolism.

In addition to nutrient substitutions and enrichment tests, we performed nutrient depletions experiments based on the Pafoba medium to identify which nutrients are required or stimulatory for *Bacillus* growth. The effect of amino acids on *Bacillus* growth was assessed using two different media variants, one medium lacking a mixture of twelve amino acids essential or conditionally essential (arginine, cysteine, tyrosine) to human nutrition, and another medium lacking a mixture of seven amino acids non-essential to humans.

Depletion of twelve (conditionally) essential amino acids had the strongest impact on *Bacillus* growth, as it significantly decreased the growth of seven strains and completely inhibited the growth of two pathogenic *Bacillus* strains. Even though we did not test the effect of depleting single essential amino acids, previous studies have established that essential amino acids valine, leucine, and threonine are required for *B. cereus* growth (23) and cereulide toxin production (57). These findings align with defined media formulations for *B. cereus*, which typically include a broad spectrum of amino acids (Table 1). The link between branched-chain amino acids and cereulide production may be explained by CodY, a key regulator of cereulide synthesis, which is activated by branched-chain amino acids acting as signals of the cell’s nutritional status (58, 59).

In contrast to *B. cereus*, we found that essential amino acids were not required for the growth of *B. subtilis, B. pumilus, B. amyloliquefaciens*, and *B. licheniformis* strains, although supplementation often stimulated growth performance. This observation aligns with previous findings that *B. subtilis* can synthesize all 20 proteogenic amino acids *de novo* (60, 61). However, some amino acid auxotrophic strains within the *B. subtilis* clade have been reported, including the wild-type *B. pumilus* DSM 18097, which requires cysteine and histidine for growth (22), as well as several mutationinduced amino acid auxotrophs (62–64). Thus, while most *B. subtilis* clade strains are prototrophic, targeted supplementation with amino acids remains valuable both for optimizing growth and for supporting strains with specific metabolic requirements.

Aside from amino acids, biotin supplementation significantly enhanced growth in five *Bacillus* strains. Both *B. pumilus* strains showed improved growth with added biotin, supporting previous recommendations to include biotin in *B. pumilus* culture media (21, 22). Interestingly, within the *B. subtilis* species, biotin’s impact on growth appeared to be strain-specific. Only the two strains isolated from food, *B. subtilis* PRO64 and PRO122 were influenced by the omission of biotin. *B. subtilis* PRO64, isolated from African fermented food, Kantong, showed almost no growth in the absence of biotin. This finding challenges earlier reports characterizing *B. subtilis* as biotin-prototrophic with a complete biosynthetic pathway (65, 66). The newly observed biotin dependency likely reflects PRO64 adaptation to a biotin-rich environment during spontaneous food fermentations, with prolonged co-cultivation with other strains which may have promoted biotin cross-feeding.

From all thirteen *Bacillus* strains tested, type strain *B. subtilis* DSM10 ^T^ was the only strain that showed stable growth in all the depletion media, while the growth of all other *Bacillus* strains was affected by at least one of the depletion media. Wang, Greenwood & Klein (2023) reported that the omission of trace metal mixture inhibited *B. subtilis* growth (42). In agreement with this, our study found that trace metals stimulated growth, although this effect was strain-dependent, observed in only two of the four *B. subtilis* strains tested. Overall, the nutrient depletion results indicate that the inclusion of vitamins, essential and non-essential amino acids, biotin, trace elements in the Pafoba medium are necessary to support the growth of a wide range of *Bacillus* strains.

The strain-specific variability observed in the stimulatory effects of certain nutrients or nutrient groups highlights the inherent diversity within the genus of *Bacillus*. The observed intraspecies variability in nutrient requirements may partly be attributed to the strains source of origin, as some were isolated from fermented food samples while others were isolated from non-food environmental niches. *Bacillus* species can in general be isolated from a wide range of environments, including soil, plant roots, fermented foods, aquatic habitats, and the gastrointestinal tracts of animals (67). Ecological origin shapes their metabolic activity and drives genetic adaptations, resulting in strain-specific nutrient requirements. Chang et al. (2020) demonstrated that *Bacillus* strains’ ability to utilize different carbon sources correlates with their ecological niches (68). This highlights the importance of including a representative variety of *Bacillus* strains in the design of a chemically defined medium, to guarantee broad applicability and strong performance across diverse *Bacillus* isolates.

All growth experiments were performed in a microtiter plate setting using an absorbance reader where mixing was only possible right before measurement. Although no visible biofilm was observed after 48 hours, the one-hour periods of static incubation between measurements may have subtly affected OD_600_ readings. Additionally, the optical density measurements could not account for the potential contribution of dead cells to growth, e.g., as result of cell lysis. Even though these factors may have influenced the OD_600_ readings, the standardized setup enabled reproducible measurements across all tested strains, facilitating comparative insights into strain-specific growth characteristics within the *Bacillus* genus.

## Conclusions

The developed chemically defined Pafoba medium proved to be an effective and versatile cultivation medium for a broad range of food-relevant and pathogenic *Bacillus* strains. All thirteen strains tested successfully grew on the Pafoba medium, out of which ten displayed a similar or higher maximum OD_600_ compared to the rich BHI medium. We systematically tested nutrient substitutions, enrichments, and depletions to identify which compounds were essential for optimal growth of *Bacillus* strains. This revealed a previously unrecognized biotin requirement for *Bacillus subtilis* PRO64, and the necessity of including essential amino acids for the growth of the pathogenic *Bacillus weihenstephanensis* and *Bacillus cereus* strains. Additionally, we identified intra-species variation in stimulatory nutrient requirements, underscoring the diverse metabolic needs among strains within the *Bacillus* genus. Overall, the Pafoba medium tested on seven *Bacillus* species, offers a controlled and reproducible experimental environment for research within the field of food safety and food fermentation.

## Acknowledgements

We would like to thank Katrine Lydom Brandenburg for assistance in the lab. The work was supported by the project PROFERMENT: Solid-state fermentations for protein transformations and palatability of plant-based foods (Grant from the Novo Nordisk Foundation under the 2021 Challenge Program, NNF21OC0066330), and the Novo Nordisk Foundation BIOBARRIER grant NNF24OC0088161. West-African strains were predominantly collected as part of the SeedFood project 09-072KU (Value Added Processing of Underutilized Savanna Tree Seeds for Improved Food Security and Income Generation in West Africa) funded by Danida. The funders had no role in study design, data collection and interpretation, or the decision to submit the work for publication.

## CRediT authorship contribution statement

Tessa S. Canoy: Conceptualization, Data curation, Investigation, Formal analysis, Methodology, Visualization, Writing – original draft, Writing – editing & reviewing; Emma S. Wiedenbein: Validation, Writing – original draft, Writing – editing & reviewing; Charlie H. McPhillips: Validation, Writing – editing & reviewing, Lene Jespersen: Funding acquisition, Supervision, Writing – editing & reviewing; Henriette L. Røder: Conceptualization, Methodology, Funding acquisition, Supervision, Writing – editing & reviewing; Dennis S. Nielsen: Conceptualization, Methodology, Funding acquisition, Project Administration, Supervision, Writing – editing & reviewing.

